# Hydrophilic/ Omniphobic droplet arrays for high-throughput and quantitative enzymology

**DOI:** 10.1101/2024.07.19.604368

**Authors:** Byungjin Lee, Fanny Sunden, Michael Miller, Bumshik Pak, Anker Krebber, Stefan Lutz, Polly Morrell Fordyce

**Affiliations:** Department of Genetics, Stanford University, Stanford, CA 94305; Department of Biochemistry, Stanford University, Stanford, CA 94305; Codexis, Inc., Redwood City, CA 94063; Department of Bioengineering, Stanford University, Stanford, CA 94305; Sarafan ChEM-H, Stanford, CA 94305; Chan Zuckerberg Biohub San Francisco, San Francisco, CA 94110

## Abstract

Engineered enzymes with enhanced or novel functions are specific catalysts with wide-ranging applications in industry and medicine. Here, we introduce Droplet Array Microfluidic Enzyme Kinetics (DA-MEK), a high-throughput enzyme screening platform that combines water-in-air droplet microarrays formed on patterned superhydrophilic/omniphobic surfaces with cell-free protein synthesis to enable cost-effective expression and quantitative kinetic characterization of enzyme variants. By printing DNA templates encoding enzyme variants onto hydrophilic spots, stamping slides to add cell-free expression mix, and imaging the resulting arrays, we demonstrate reproducible expression of hundreds of enzyme variants per slide. Subsequent stamping of fluorogenic substrates and time-lapse imaging allows determination of Michaelis-Menten parameters for each variant, with measured catalytic efficiencies spanning 5 orders of magnitude and agreeing well with values obtained via traditional microtiter plate assays. DA-MEK consumes orders of magnitude less reagents than plate-based assays while providing accurate and detailed kinetic information for both beneficial and deleterious mutations. In future work, we anticipate DA-MEK will provide a powerful and versatile platform to accelerate enzyme engineering and enable screening of large variant libraries under diverse conditions.

## Introduction

Enzymes are the catalytic machines that drive chemical reactions. Thus, they have enormous value and potential in industry (*e*.*g*. as engineered lipases and proteases for detergents, novel catalysts for bioremediation and green chemistry, and degradative enzymes for cellulosic biofuels^1-10^) and medicine (*e*.*g*. as targets for drugs used to treat cancer^11-16^ and infectious disease^17-21^, as catalysts for efficient synthesis of pharmaceuticals^22-25^, and as therapeutics in metabolic flux disorders^26-29^). Despite impressive and rapid progress in the ability to design protein sequences that reliably adopt desired low-energy structures, designing enzymes with specified activities, specificities, and catalytic efficiencies remains an unsolved challenge. As a result, engineering enzymes with enhanced or novel functions typically involves multiple rounds of experimental directed evolution.

Directed evolution campaigns are often costly and labor-intensive, as each round necessitates the synthesis of variant DNA libraries, expression of variant proteins, and functional characterization of enzymatic activity. At industrial scale, these steps often take place within 384-well plates that require approximately 10-20 µL of material for each reaction at each step. Despite these requirements, advancements in microfluidic technologies present innovative solutions to overcome these limitations. Microfluidic droplet assays enable ultra-high-throughput compartmentalization of reactions within very small volumes (*e*.*g*. ∼ 1,000,000 droplets/hr and 6-pL/droplet), reducing reaction costs by up to 10^6^-fold^30, 31^. These droplets can be coupled to fluorescence- or absorbance-based assays and different sorting technologies (*e*.*g*. FADS^30-35^ or FACS^36-39^) to select variants with desired activities. However, the enrichment of only variants with enhanced activities precludes gathering information about deleterious sequence changes that can be critical for building a protein design tool to predict activities from sequences. Furthermore, these approaches cannot directly link measured activity of individual variants with their sequences.

To address the need for new technologies capable of providing quantitative functional characterization of many enzyme variants in parallel, our lab previously developed HT-MEK (High-Throughput Microfluidic Enzyme Kinetics), which relies on valved microfluidic devices aligned to spatially arrayed expression plasmids to recombinantly express, purify, and functionally characterize up to 1,800 enzyme variants in parallel. HT-MEK returns quantitative thermodynamic and kinetic constants (*e*.*g. K*_i_, *k*_cat_, *K*_M_, and *k*_cat_/*K*_M_) for each variant in the presence of multiple substrates and inhibitors and under different conditions, thus providing both ‘positive’ and ‘negative’ information about sequence changes that either improve or reduce activities. However, implementing HT-MEK requires fabrication of complex two-layer PDMS valved microfluidic devices and significant custom hardware for on-chip pneumatic valve control,^40^ posing technological barriers to widespread adoption.

Droplet microarrays on superhydrophobic/superhydrophilic or omniphobic/superhydrophilic surfaces provide a promising and easily adoptable method for similar array-based droplet compartmentalization. Patterned surfaces comprised of superhydrophilic spots surrounded by superhydrophobic regions create virtual barriers to aqueous fluids, enabling spontaneous formation of µL-to-nL water-in-air droplet arrays and multiplexed compartmentalization. The open format of these arrays facilitates easy reagent addition or sampling by ‘stamping’ slides vertically together^41, 42^. Demonstrating their utility and flexibility, superhydrophobic/superhydrophilic droplet arrays have already been used for a variety of high-throughput screening applications, including synthesizing chemicals, screening cells and drugs, and designing materials^43-59^. To date, however, such wettability-patterned droplet arrays have not been used for *in vitro* enzyme expression and downstream characterization of the expressed enzymes.

Here, we demonstrate high-throughput cell-free expression and functional characterization of enzyme catalytic efficiency (DA-MEK, for <u>D</u>roplet <u>A</u>rray <u>M</u>icrofluidic <u>E</u>nzyme <u>K</u>inetics) within omniphobic/superhydrophilic droplet arrays using variants of PafA, a well-characterized model alkaline phosphatase. By printing DNA plasmids encoding the expression of C-terminally eGFP-tagged PafA variants onto hydrophilic spots, ‘stamping’ these spots to a matched array containing cell-free protein expression mixture, incubating to produce protein, and imaging to quantify protein expression, we establish the ability to reproducible express >500 nM protein per droplet. By subsequently ‘stamping’ these expressed protein arrays to a matched array containing a fluorogenic PafA substrate and imaging to quantify turnover over time, we establish the ability to quantify reaction turnover and determine Michaelis-Menten rate constants for the variants of the enzyme in parallel. Rate constants measured via DA-MEK agree well with values derived from traditional plate-based assays (r^2^= 0.89 and RMSE = 0.2) for PafA variants spanning a catalytic efficiency range of over 5 orders of magnitude.

## Results

### Pipeline for high-throughput recombinant enzyme expression and kinetic characterization within droplet arrays

To collect enzyme kinetics data, we print a library of DNA templates encoding expression of C-terminally eGFP-tagged variants onto hydrophilic spots patterned within a larger omniphobic surface (either manually or using a robotic printer, see **Methods**). To express protein variants, we ‘stamp’ a second patterned slide containing reagents for cell-free protein expression (CFPS) onto this printed library slide to add CFPS reagents to each spot containing a DNA template. Following the addition of CFPS reagents, slides are incubated for 4 hours at 23°C, allowing each droplet to act as a miniaturized protein-producing bioreactor. Expressed protein can be quantified via eGFP-associated fluorescence, allowing for simultaneous, non-contact quantification of protein expression over time and across variants. To quantify reaction kinetics for each variant in parallel, we prepare a third patterned slide with fluorogenic substrate onto the slide containing expressed protein to start enzymatic reactions and then image over time. Quantifying substrate fluorescence over time makes it possible to quantify reaction progress over time and ultimately determine functional kinetic parameters (*e*.*g. k*_cat_, K_M_, and *k*_cat_/K_M_) (**Figure 1A**).

**Figure 1.**
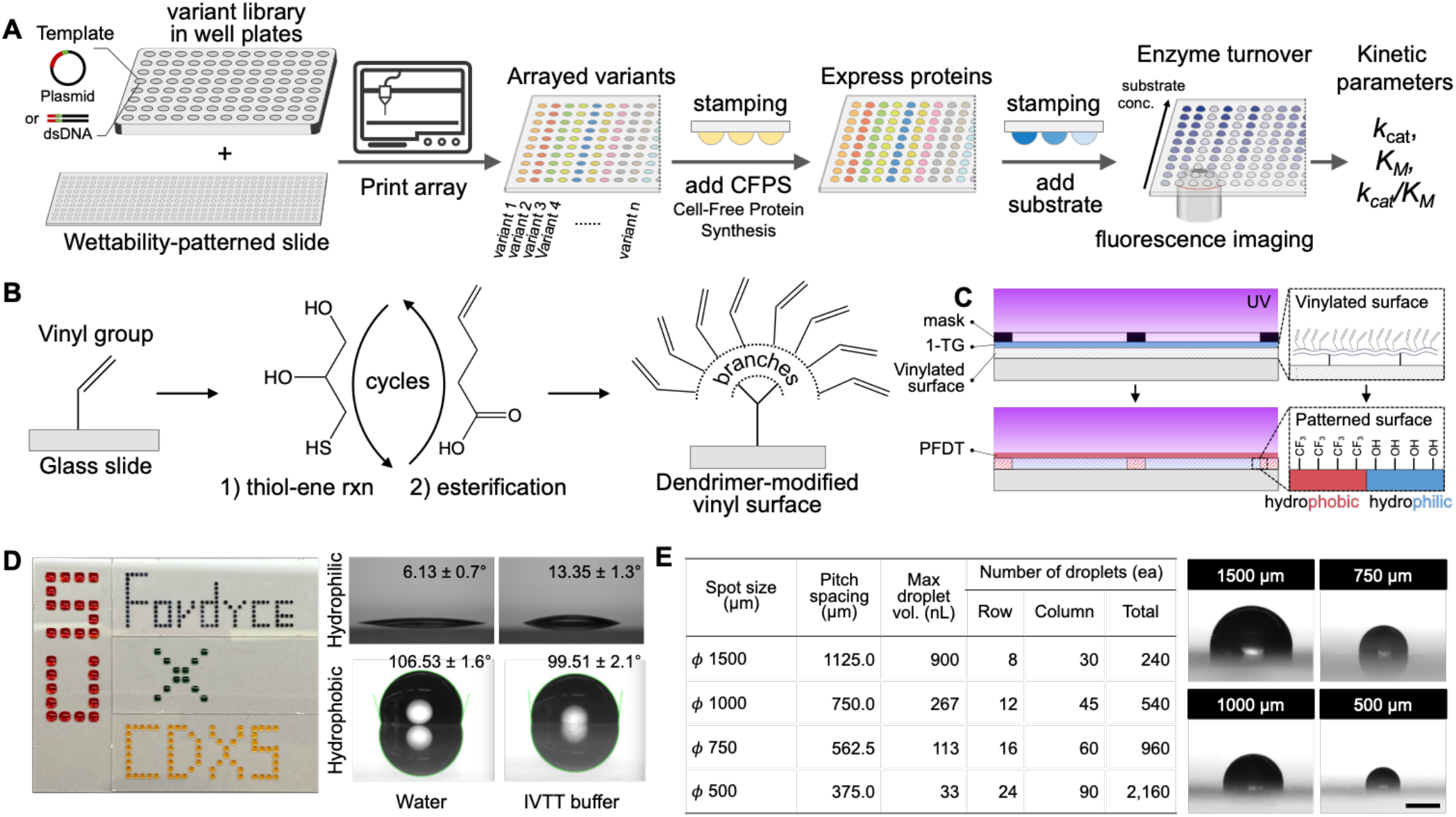
Workflow for Droplet Array Microfluidic Enzyme Kinetics (DA-MEK). **A)** Experimental pipeline: (1) DNA templates encoding enzyme variants are printed onto wettability-patterned slides, (2) arrayed variants are ‘stamped’ to another patterned slide to add cell-free protein expression (CFPS) mix to each variant and express proteins, and then (3) expressed proteins are ‘stamped’ to fluorogenic substrates to begin reactions. Imaging over time allows quantification of protein expression and substrate turnover for all variants in parallel. **B**) Chemical steps for modifying the glass surface to introduce a dendrimer-modified vinyl surface via thiol-ene reactions with 1-Thioglycerol (1-TG) and esterification with 4-Pentenoic acid. **C**) Patterning surface wettability using UV light and a mask through thiol-ene photoclick chemistry with 1-Thioglycerol (1-TG) and perfluorodecanethiol (PFDT). **D)** Examples of droplet array formation on slides with different spot sizes (left); images and contact angles of water and CFPS reagents on hydrophilic and hydrophobic surfaces (right). **E)** Table summarizing droplet array geometries and volumes for different droplet sizes (left) and bright field images of droplets on hydrophilic spots of different diameters (right); scale bar indicates 500 μm.

### Patterned slide surfaces drive robust formation of uniform water-in-air droplet arrays

To fabricate droplet microarray (DMA) slides, we adopted a surface-tethered dendrimer modification scheme^41^ that proceeds via thiol-ene photoclick reactions and Steglich esterification^41, 60^ (**Figure 1B**). First, 1-thioglycerol is covalently coupled to vinylsilanes attached to the glass surface via exposure to UV light, leading to the formation of two hydroxyl groups; subsequent esterification with 4-pentenoic acid reintroduces vinyl groups to each hydroxyl. Repeating this process three times expands a single vinyl group to eight vinyl groups, simultaneously increasing the density of chemical moieties and enhancing the surface area of the smooth glass to yield improved hydrophilic and omniphobic properties^41^. Coupling 1-thioglycerol to the slide surface and covering it with a designed photomask that only allows light to penetrate in circular spots across the slide surface yields an array of hydrophilic spots **(Figures 1C and S1)**; subsequently treating the remaining area with 1H,1H,2H,2H-perfluorodecanethiol creates omniphobic barriers with a low surface free energy surrounding each hydrophilic spot. This results in the formation of a virtual well plate consisting of invisible hydrophilic patches and omniphobic walls (**Figure 1D**, left), facilitating the organization and segregation of different chemical or biological samples.

Aqueous liquids deposited onto hydrophilic and omniphobic surfaces spread out or round up to form droplets, respectively, with the contact angle of the droplet determined by the relative hydrophilicity or hydrophobicity of the surface and properties of the aqueous fluid. Measured contact angles for deionized water pipetted onto hydrophilic and omniphobic regions were 6.1 ± 0.7° and 106.53 ± 1.6°, respectively; the difference between contact angles for cell-free protein expression product was slightly lower (13.4 ± 1.3° and 99.5 ± 2.1° on hydrophilic and omniphobic regions, respectively), which is thought to be due to the adsorption of the expressed protein in the cell-free protein expression product onto the surface or the reduced surface tension of the droplet caused by the expressed protein (**Figure. 1D**, right). Water-in-air droplets remained well-separated over time, enabling the creation of arrays with varying numbers of droplets containing different fluid volumes (**Figure 1E**). To determine the maximum droplet volume that can be accommodated on each hydrophilic spot, we empirically determined the largest volume at which the droplet contact angle stabilized at 100° or less for a 1500 µm droplet and used this as a reference point to estimate the other volumes for the various spot sizes (**Figure 1E**).

### Water-in-air droplet arrays enable reproducible high-throughput recombinant protein expression

Prior work has established the ability to add reagents to or draw reagents from water-in-air droplet arrays via a ‘stamping’ process in which two arrays are brought in close proximity vertically such that droplets merge while hydrophobic forces prevent adjacent droplets from mixing^41, 42^. Here, we leverage this principle for high-throughput protein expression by: (1) printing plasmids encoding expression of protein variants on one DMA slide, (2) preparing a second DMA slide with droplets of identical spacing containing reagents required for cell-free protein expression with an automated dispenser (**Figure S2**), (3) loading DMA slides into a custom-built micro-manipulatable aligner to bring them together to mix reagents, and then (4) incubating to allow protein expression (**Figures 2A and B**).

**Figure 2.**
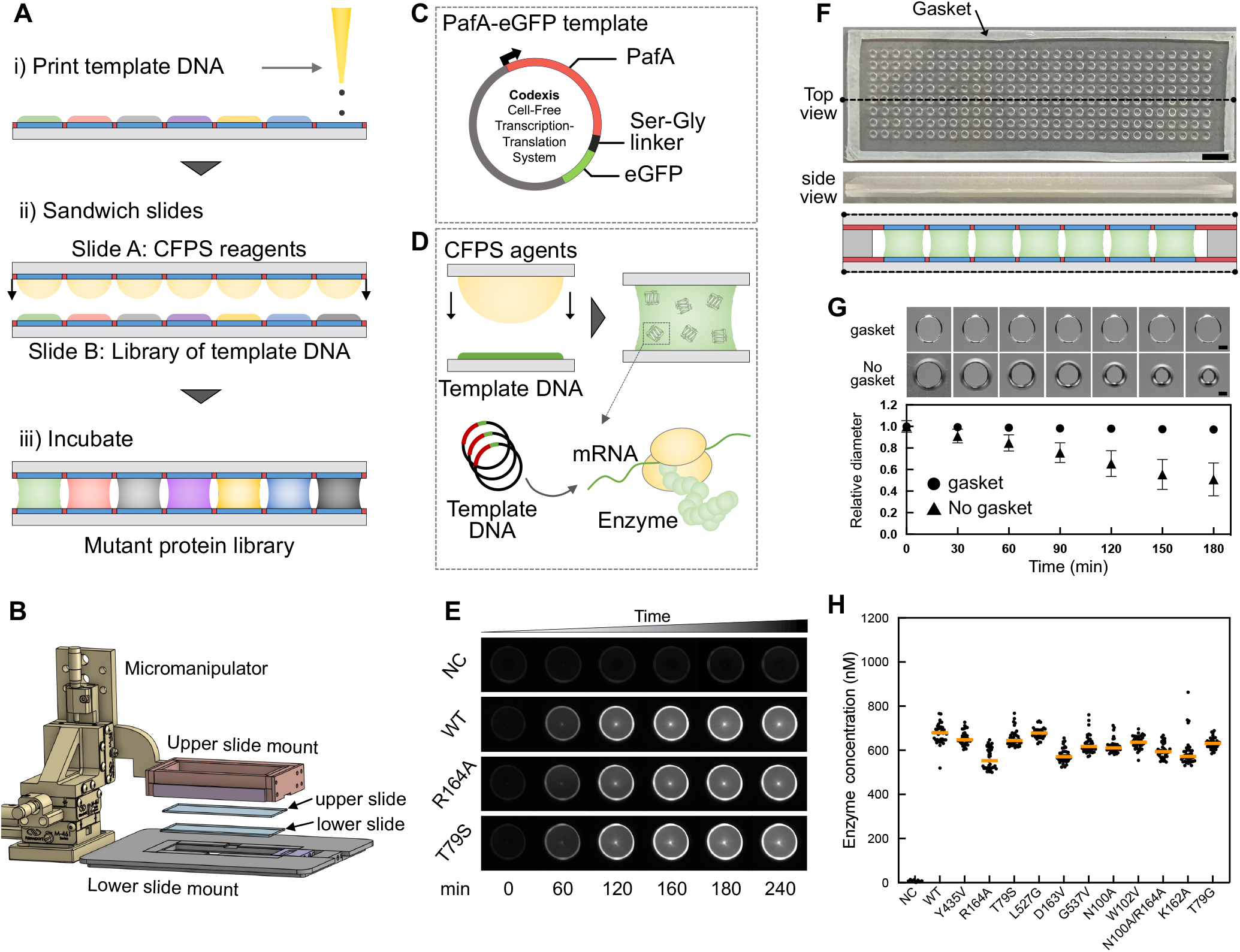
Cell-free protein expression by stamping droplet microarrays. **A)** Experimental pipeline for cell-free protein expression via (i) printing template DNA encoding enzyme variants onto hydrophilic spots, (ii) stamping this template DNA DMA slide with a DMA slide containing cell-free protein expression mix in each droplet, and (iii) incubating to express the mutant protein library. **B)** A 3D rendering of microscope stage-mountable slide aligner to ‘stamp’ DMA slides together. **C)** Diagram illustrating the plasmid DNA template used to produce C-terminally eGFP-tagged PafA variants. **D)** Diagram illustrating how addition of cell-free protein synthesis reagents to template DNA yields mRNA and expressed eGFP-tagged protein. **E)** Fluorescence microscopy images illustrating cell-free expression of PafA-eGFP variants at various time points. **F)** Bright field microscopy images and schematic showing gasket used to minimize droplet evaporation. Scale bar indicates 5 mm. **G)** Bright field microscopy images (top) and quantified droplet diameters (bottom) for droplets over time with and without the gasket. Scale bars indicate 500 μm. **H)** Calibrated enzyme concentrations produced for droplets expressing 12 different eGFP-tagged PafA variants across 2 different DMA slides.

To evaluate DMA protein expression, we generated 12 plasmid templates encoding C-terminally eGFP-tagged variants of the phosphate monoesterase PafA, a model enzyme with previously characterized active site mutants whose catalytic efficiencies span >5 orders of magnitude ^61, 62^ (**Figure 2C and D**). We ‘stamped’ slides bearing these templates with slides containing cell-free expression mixtures and then quantified eGFP fluorescence over time via microscopy to estimate expressed protein concentrations (**Figure 2E and S4**). Initial trials of protein expression revealed significant evaporation and droplet shrinkage during the 3-hour required incubation at 23°C, altering buffer concentrations and reducing protein expression (**Figure 2G**). To ameliorate this, we designed a gasket that sits between the two glass slides to prevent droplet evaporation during incubations or enzyme turnover processes, serving as both a sealant and a spacer (**Figure 2F**). The gasket was made via hot embossing of multiple layers of Parafilm and is cut to occupy approximately 2 mm along the outline of the slides (25 × 75 mm). Before gasket addition, droplets showed a 50% reduction in diameter during a 3-hour incubation at 23°C; after gasket addition, droplet diameters remained unchanged over the same timeframe (**Figure 2G**).

Measured intensities for droplets containing printed plasmids encoding expression of either wild-type PafA or single-nucleotide variants showed steady increases in fluorescence over time while negative control droplets lacking plasmids remained dark, as expected (**Figure 2E**). To quantify protein yields, we generated calibration curves by adding the same volumes of purified PafA-eGFP (86.4 kDa) at known concentrations and quantifying intensity (**Figure S3**). Calibrated protein intensities across multiple droplets on a given slide showed strongly reproducible variant expression with final concentrations ranging between 500-800 nM protein in 900-nL droplet volumes (38.9-62.2 ng per variant) (**Figure 2H**). Measured fluorescence intensities were uncorrelated with previously determined catalytic efficiencies for each variant, confirming that phosphate monoesterase activity does not impact expression (**Figure S5**). Protein expression was relatively constant across different variants and did not correlate with either droplet position within a slide or experimental replicate (**Figure 2H** and **Figure S6**).

### Water-in-air droplet arrays enable quantitative kinetic measurements of enzymatic turnover with a wide dynamic range within low volumes

Next, we tested whether DMA stamping could enable high-throughput and quantitative measurements of enzyme catalysis by: (1) merging a DMA array containing expressed enzyme variants with a second array containing fluorogenic substrate, and then (2) imaging over time to detect the production of a fluorescent product (**Figure 3A**). To monitor PafA phosphatase activity, we employed a coumarin-based fluorogenic 4-Methylumbelliferyl Phosphate (4-MUP) substrate that yields a fluorescent 4-Methylumbelliferone (4-MU) product upon hydrolysis (**Figure 3B and Equation S1**). Fluorescent images of PafA-variant-containing droplet arrays after stamping to fluorogenic 4-MUP substrates showed variant-dependent differences in measured fluorescence intensities over time consistent with enzymatic turnover (**Figure 3C**). As expected, wild-type PafA yielded the highest enzymatic activity; the R164A and T79S mutant constructs showed slightly and dramatically lower enzymatic activities, respectively. Negative control droplets containing all of the reagents required for cell-free protein synthesis except for plasmid template showed no increased fluorescence with time, establishing little to no background hydrolysis in the absence of expressed enzyme (**Figure 3C**).

**Figure 3:**
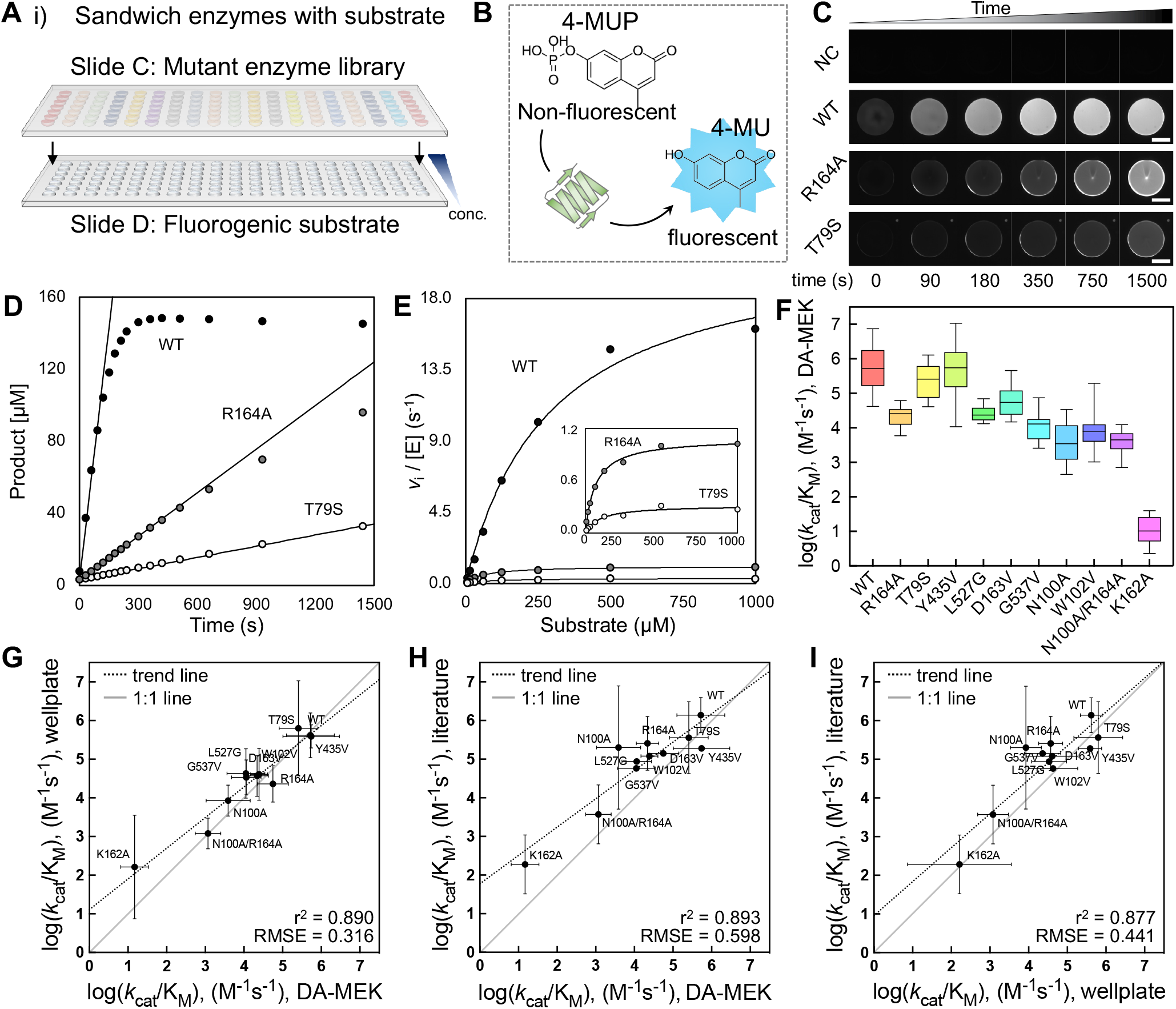
Quantifying variant enzyme activity and kinetics by stamping. **A)** Experimental pipeline for quantifying enzyme activity via stamping and expressed DMA array to an array containing fluorogenic substrate. **B)** Schematic of fluorogenic reaction in which enzymatic removal of a phosphate group from a non-fluorescent substrate (4-MUP) generates a fluorescent product (4-MU). **C)** Time-lapse images showing increased fluorescence over time for 4 example droplets either lacking DNA template (negative control, NC) or containing 3 different PafA constructs (WT, R164A, and T79). **D)** Quantified fluorescence over time for the same droplets containing either wild-type PafA (WT) or two mutant enzymes (R164A and T79S). Markers indicate median pixel intensity for a single droplet; lines indicate linear fits to initial rates. **E)** Observed initial rates as a function of substrate concentration for WT, R164A, and T79S. Markers indicate single droplets; lines indicate Michaelis-Menten fits to the data. **F)** Box plots showing measured corrected log(*k*_*cat*_/K_M_) for multiple enzyme variants obtained in DA-MEK. **G)** Corrected log(*k*_*cat*_/K_M_) values from DA-MEK versus those from wellplates. **H)** Scatter plot comparing observed log(*k*_*cat*_/K_M_) values from DA-MEK versus prior literature results. **I)** Corrected log(*k*_*cat*_/K_M_) values from DA-MEK versus prior literature results. The data across figures F to I are based on 15 measurements for each variant.

Using calibration curves **(Figure S7)** to convert measured intensities to the amount of product generated allows quantitative estimation of substrate turnover over time within each droplet (**Figures S10 and S13**); plotting initial rates as a function of substrate concentration and fitting to the Michaelis-Menten equation then allows estimation of kinetic parameters (*k*_cat_, *K*_M_, and *k*_cat_/*K*_M_) **(Figures S11 and S14)**. As cell-free expression mixtures contain substantial inorganic phosphate (a known ground state inhibitor for PafA), we: (1) quantified the amount of inorganic phosphate within cell-free expression mixtures using a standard malachite green assay (**Figure S8**), and then (2) corrected both the initial velocities at each substrate concentration and the fitted K_M_ using competitive inhibition model equations (**Equation S2, S3, and Figure S9** in Supplementary Information). For each mutant, we used previously-measured inhibition constants for each variant and measured inorganic phosphate levels to yield accurate values^61, 62^. Measured *k*_cat_/*K*_M_ values were highly reproducible across mutants and spanned nearly 5 orders of magnitude (**Figures 3F, S12, and S15**), with the lowest resolvable activity of ∼ 14.6 M^-1^s^-1^ for a catalytically compromised K162A variant.

### Water-in-air droplet array measurements recapitulate traditional plate-based values

To assess the accuracy and precision of DA-MEK catalytic activity measurements, we measured catalytic activities for the same set of variants via a traditional plate-based assay using the same cell-free expression mix and directly compared values obtained via DA-MEK and plate-based assays (**Figures 3G-I and S16**). DA-MEK measurements of *k*_cat_/*K*_M_ agreed extremely well with plate-based measurements (*r*^2^ = 0.89 and RMSE = 0.20) (**Figure 3G**), establishing the ability to acquire high-quality estimates of catalytic activity while consuming 11-fold fewer reagents per reaction. Although DA-MEK experiments quantified turnover for enzymes in solution and in the presence of cell-free expression mix while prior HT-MEK experiments quantified turnover for surface-immobilized enzymes following washing and purification, the results from both techniques also showed strong agreement with one another (*r*^2^ = 0.893 and RMSE = 0.663), as shown in **Figure 3H**. The largest measured disagreement was for PafA N100A (3.37 vs 4.89); as N100A is strongly inhibited by inorganic phosphate ions, this discrepancy likely results from uncertainty in measurements of the amount of background inorganic phosphate within the DA-MEK arrays following cell-free protein expression.

## Discussion

Protein production and characterization are the costliest part of directed evolution campaigns. As this process often requires large volumes of expensive reagents, reduced reaction volumes directly translate into significant cost savings. Here, we demonstrate a miniaturized screening platform, DA-MEK, that enables parallelized production and functional characterization of enzymes within nL-to-µL volume droplets. Compared to conventional 384 well plate-based assays, this reduces per-reaction costs by approximately 11-fold for the 450 nL droplets used here for protein synthesis and enzymatic assays (**Table S1**). As libraries grow in size, these savings can be substantial. Assuming 8 substrate concentrations per Michaelis-Menten curve and 3 replicates per measurement, characterizing a 1000-member library costs $12,744 using DA-MEK with 1500 µm diameter spots as compared to $141,600 for 384-well plates (**Table S1**). Future experiments optimizing protein synthesis and enzymatic turnover measurements in 500 µm diameter spots could increase these savings further, reducing estimated costs for the same 1000-member library to $566, 250-fold less than 384-well plates.

These savings are a conservative estimate that do not consider additional savings associated with the fact that DA-MEK does not require pipette tips for reagent addition.

To date, water-in-oil single- and double-emulsion droplet-based assays have had limited application to high-throughput pharmaceutical screening, as the low molecular weight and hydrophobicity associated with candidate drug molecules cause them to diffuse out of aqueous compartments and into the continuous oil phase. Separating droplet compartments with an air barrier via droplet self-assembly on superhydrophilic/superhydrophobic surfaces eliminates this potential source of cross-contamination. In future work, DA-MEK protein arrays could be ‘stamped’ directly to patterned slides bearing small molecules to functionally screen for drug inhibition of enzymatic activity in high-throughput; iterative stamping of different slides could facilitate combinatorial drug screening. As extensive prior work has established that superhydrophilic/superhydrophobic droplet arrays can be compatible with label-free readouts like mass spectrometry^43, 44, 52, 55, 63, 64^, DA-MEK could be coupled with a mass spectrometry readout to screen for enzymatic activities that are not easily profiled via fluorescence. A mass spectrometry readout would also be highly useful for commercially relevant directed evolution campaigns, as assaying the true substrate (and not a dye-labeled version) is essential to ensure that evolution targets the real compound rather than merely the model compound. Finally, DA-MEK’s use of cell-free protein synthesis enables future screening of proteins toxic to cells (e.g. many nucleases and proteases), facilitates activity measurements in the presence of chemical denaturants (e.g. urea or guanidinium chloride) to disambiguate folding and catalytic effects, and is compatible with recently-developed genetic code expansion methods that enable incorporation of non-canonical amino acids^65-67^.

As industrial-scale directed evolution efforts generate large data sets, random mutagenesis has begun to be replaced by machine-learning-guided sequence design. For example, a recent study demonstrated the potential of droplet microfluidics for high-throughput profiling of a biocatalyst, an imine reductase (IRED), for training a machine-learning protein design model by screening a library of over 17,000 enzyme variants^68^. While most droplet-based screening approaches provide ‘positive’ information about variants with desired activities, they cannot return ‘negative’ information about deleterious substitutions that reduce activity that is particularly valuable for accurate model training^62, 69, 70^. DA-MEK’s array-based format offers direct measurement of turnover for all variants (and not just those recovered during selection), providing valuable information about sequence changes that both enhance and reduce observed activities. Finally, DA-MEK’s open format is highly amenable to automation. Thus, we anticipate that DA-MEK could provide a scalable method for efficient high-throughput screening of large variant libraries within directed evolution and drug screening campaigns.

## Materials and Methods

### Photomask design and production

Photomasks were designed using AutoCAD software (Autodesk, Inc.). Hydrophilic spots were designed to be transparent with diameters of 1500, 1000, 750, and 500 µm, and the spacing between adjacent spots was set to half the diameter of each spot. These designs were produced on a 5-inch chrome quartz mask (Front Range Photomask, Las Vegas, NV, USA). Details and specifications of all mask files are available for download in the OSF repository associated with the paper (osf.io/aqtzr).

### Surface patterning

We activated Nexterion glass B slides (25 × 75 mm, Schott) using Piranha cleaning (H_2_SO_4_:H_2_O_2_ = 1:3) and then treated them with a three-cycle series of chemical reactions for dendrimer generation^41^: For thiol-ene photoclick chemistry, we applied a mixture of 1 wt% 2,2-Dimethoxy-2-phenylacetophenone (cat. no. 196118, Sigma-Aldrich) and 10 wt% 1-Thioglycerol (cat. no. 88640, Sigma-Aldrich) in DI water/ethanol (1:1 vol) to slide surfaces, covered them with a fluorinated quartz plate, and then exposed this assembly to 254 nm UV light (36W, VirusLights, Delano, TN, USA) for 3 minutes at 3 cm distance. After exposure, we washed slides with ethanol and then dried them using nitrogen gas and an 80°C convection oven for 5 min. For esterification, we immersed the dried slides in an esterification mixture consisting of 200 mg 4-(dimethylamino)pyridine (cat. no. W284300, Sigma-Aldrich), 444 μL pentenoic acid (cat. no. 8204990005, Sigma-Aldrich), and 180 μL N,N’-Diisopropylcarbodiimide (cat. no. 8036490010, Sigma-Aldrich) in 200 mL of acetone for 4 hours. After the final cycle, we initiated the thiol-ene reaction using a quartz chrome mask with clear spots (**Fig. S1**) for hydrophilic spot patterning. To complete the functionalization, we then washed and dried each slide, coated them with a 10% 1H,1H,2H,2H-perfluorodecanethiol (PFDT) (cat. no. 660493, Sigma-Aldrich) solution in acetone, covered with a fluorinated blank quartz plate, and again exposed to 254nm UV light (36 W at 3 cm distance for 3 min) to complete the functionalization of the surrounding area^41^.

### Contact angle measurement

We measured contact angles using a goniometer (Model 290, ramé-hart instrument, Succasunna, NJ, USA). 5 µL droplets of two different liquids (deionized water or cell-free protein expression (CFPS) reagent) were carefully applied onto hydrophilic or omniphobic-treated glass slides, respectively. For each type of treatment and liquid, images of 15 droplets were captured and analyzed using an ImageJ plugin (DropAnalysis^71^).

### Gasket design and assembly

To create a gasket that could prevent droplet evaporation during protein expression, we employed one of two methods: (1) hot embossing four layers of Parafilm at 80°C or (2) pouring and curing a mixture of polydimethylsiloxane (PDMS) (RTV-615, Momentive) in a 10:1 ratio between two fluorinated glass plates. We used pieces of Whatman Thin Layer Chromatography Plates (cat. no. 4861-830, Thickness: 500 µm) as spacers. The thickness of the final product was inspected and then trimmed to create a rectangular window measuring 21 × 71 mm, ensuring a 2 mm overlap on the 25 × 75 mm slides.

### Design and fabrication of Aligner

To accurately align and ‘stamp’ 2 DMA slides together, we designed a manual aligner that includes a lower holder in the form of a microscope-mountable insert and an upper holder with a feature for magnetic attachment and detachment to a 3-axis micromanipulator (Newport M-461). These components were fabricated using CNC machining and subsequently assembled and mounted to a motorized *xy-* stage (MS-2000, Applied Scientific Instrumentation) on a Nikon Ti-2 automated fluorescence microscope. All design files required to produce the aligner are available within the OSF repository associated with the manuscript (osf.io/aqtzr).

### Printing of plasmid libraries onto droplet array slides

Plasmids encoding expression of C-terminally eGFP-tagged PafA variants were produced via Gibson assembly and QuickChange mutagenesis (Agilent) as previously described^62^. To print plasmids onto the slides, we used a SciFlexArrayer S3 (Scienion) equipped with a PDC70 piezoelectric capillary nozzle (P2030-S6041, Scienion) to print 150 nL of plasmid DNA (at a concentration of 100-300 ng/µL) in deionized water (DIW) (∼100 ng/ml) onto each hydrophilic spot of the slide. Between the printing of each variant, the nozzle was washed three times and dried for 15 seconds in an ambient environment.

### Cell-free protein expression

To produce protein, we used either cell-free expression mix provided by Codexis (Redwood City, CA, USA) or the PURExpress kit from New England Biolabs (USA). Mixtures were prepared according to the manufacturer’s protocol with the addition of RNase inhibitor and then loaded onto DMA slides using a custom-built automated liquid dispenser (**Figure S2**). After stamping slides to add cell-free protein expression mix to printed DNA templates, we incubated at 23°C for 4 hours to express proteins (unless otherwise stated).

### Microscopy setup

All images were obtained using a Nikon Ti-S Microscope which includes a motorized XY stage from Applied Scientific Instrumentation (MS-2000 XY stage), a CMOS camera by Oxford Instruments (Andor Zyla 4.2), a solid-state light source by Lumencor (SOLA SE Light Engine), and an automated filter turret. This turret was outfitted with eGFP and DAPI filter sets from Chroma Technology Corp. (part no. 49002) and Semrock Inc. (catalog no. DAPI-1160B-NTE), respectively. Imaging was carried out with a 2× objective lens (CFI Plan Apo l 2× NA 0.10, Nikon) using 2×2 binning settings (resulting in an image resolution of 1024×1024 pixels). The exposure times were set to 500 ms for eGFP, with DAPI exposures at 50 ms.

### Generation of eGFP and 4-MU calibration curves

The concentration of eGFP produced in CFPS buffer was determined by comparing its absorbance at 488 nm to that of a blank CFPS reagent, utilizing a molar extinction coefficient of 56,000 M^-1^cm^-1^. This measurement was conducted using a UV-vis spectrophotometer (DeNovix, Wilmington, DE, USA). To correlate the fluorescent signal intensities of eGFP product observed under the microscope with the measured eGFP concentration, we imaged the eGFP product droplets at 2x magnification, droplet heights of 500 μm, and exposure times of 500 ms. We then performed a linear fit of measurements of observed intensity *vs*. known concentration for each condition and used these linear fit parameters to estimate eGFP concentrations from images of expressed protein. For the calibration of 4-methylumbelliferone (4-MU, Sigma-Aldrich), we prepared solutions of 4-MU in concentrations of 1000, 500, 250, 125, 62.5, 31.25, 15.6, and 7.8 μM, imaged to quantify droplet intensities, and again performed a linear fit of observed intensities *vs*. known concentrations. To estimate substrate turnover, we used the linear fit parameters to estimate concentrations from observed fluorescence. The representative fluorescence signal intensity for each droplet was determined by calculating the median signal intensity measured within a circular region that has a diameter 0.7 times that of the droplet.

### Enzyme kinetics measurements

4-MUP was loaded onto the substrate slide at varying concentrations (2000, 1000, 500, 250, 125, 62.5, 31.25, and 15.63 μM), each mixed with an equal volume of enzyme droplets to achieve halved concentrations. On a separate slide with the expressed enzyme, dilutions ranging from 8 to 32 times were performed using a merging-and-splitting process with a reaction buffer (100 mM MOPS, 500 mM NaCl, 100 μM ZnCl_2_, pH 8.0) slide. Enzyme and substrate slides were then placed on the microscope stage. The aligner was used to facilitate the merging of droplets from both slides. Time-lapse imaging was conducted in 30-second intervals for one hour.

### Image processing and intensity quantification

Images were analyzed using a custom Python script. Briefly, images captured at each time point were stitched together to form a single composite image. After stitching, we identified outlines and centroids of droplets captured in bright field images, defined a region of interest (ROI) for each droplet (consisting of a circle with a diameter equaling 0.7 times that of the droplet, centered on the droplet’s centroid), and the computed the median intensity within each ROI at each timepoint.

### Data analysis to estimate rate constants

A custom Python script was used to analyze the initial velocity of the process curve for each droplet. The initial velocity was determined by calculating the slope of the linear region of the curve up to the maximum time that the goodness of fit of the linear slope remained equal to or above 0.97. The parameters of the Michaelis-Menten model, *k*_cat_ and K_*M*_, were estimated using the curve_fit function from the SciPy library with initial estimates based on the substrate concentration and reaction rate data from experiments.

### Plate-based assays to quantify enzymatic activity

Concentrations of proteins expressed in tubes, using either Codexis CFPS mix or PURExpress, were initially quantified using a spectrophotometer. Subsequently, these protein solutions were appropriately diluted to become 2x the final concentration. These diluted solutions were dispensed into a 384-well plate, allocating 10 μL to each well. 4-MUP solutions were prepared at various concentrations (2000, 1000, 500, 250, 125, 62.5, 31.25, and 15.63 μM) and introduced into the enzyme-containing wells using a multi-channel pipette. A Tecan Infinite M Plex spectrophotometer (Männedorf, Switzerland) was employed to measure fluorescent intensities every 30 seconds for one hour at Ex355/Em460.

### Malachite green assay for quantifying inorganic phosphate concentration

The malachite green assay kit (MAK307, Sigma-Aldrich) was purchased and used according to the manufacturer’s protocol. A series of phosphate standard solutions with final concentrations of 40, 32, 24, 16, 12, 8, 4, and 0 μM, containing malachite green, were prepared, and their absorbance was measured at 644 nm. The protein crude product, prepared using the previously described recipe, was diluted to fall within the appropriate measurement range, and its absorbance was measured. The measured values were used to determine the concentrations through a standard curve and then the original concentrations were obtained by multiplying the dilution factors.

