## Supplemental Information for "Hydrophilic/ Omniphobic droplet arrays for high-throughput and quantitative enzymology"

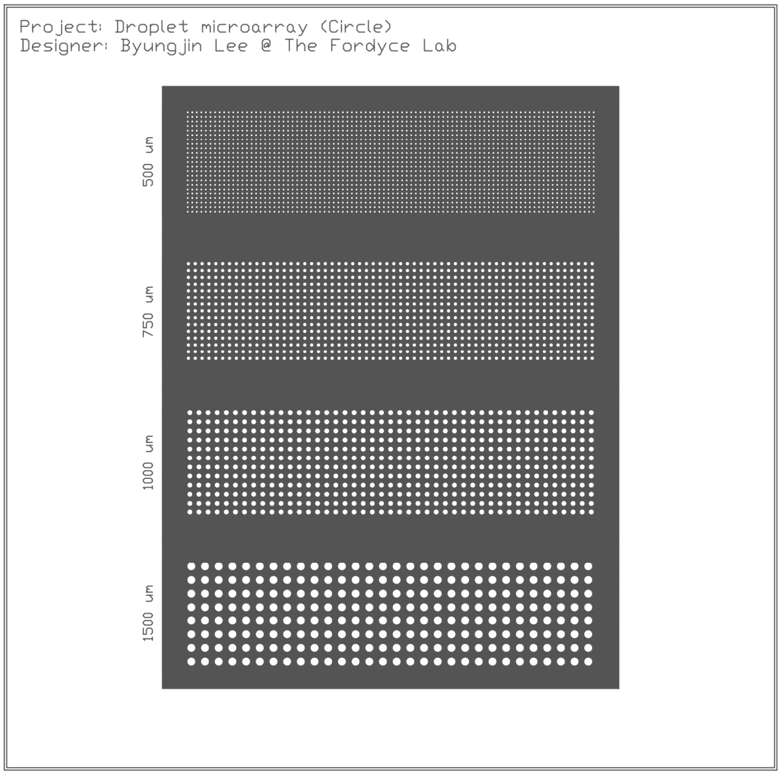


**Figure S1.** Design of photomask for thiol-ene photoclick chemistry. Mask files are available for download in the OSF repository associated with the paper (osf.io/aqtzr).

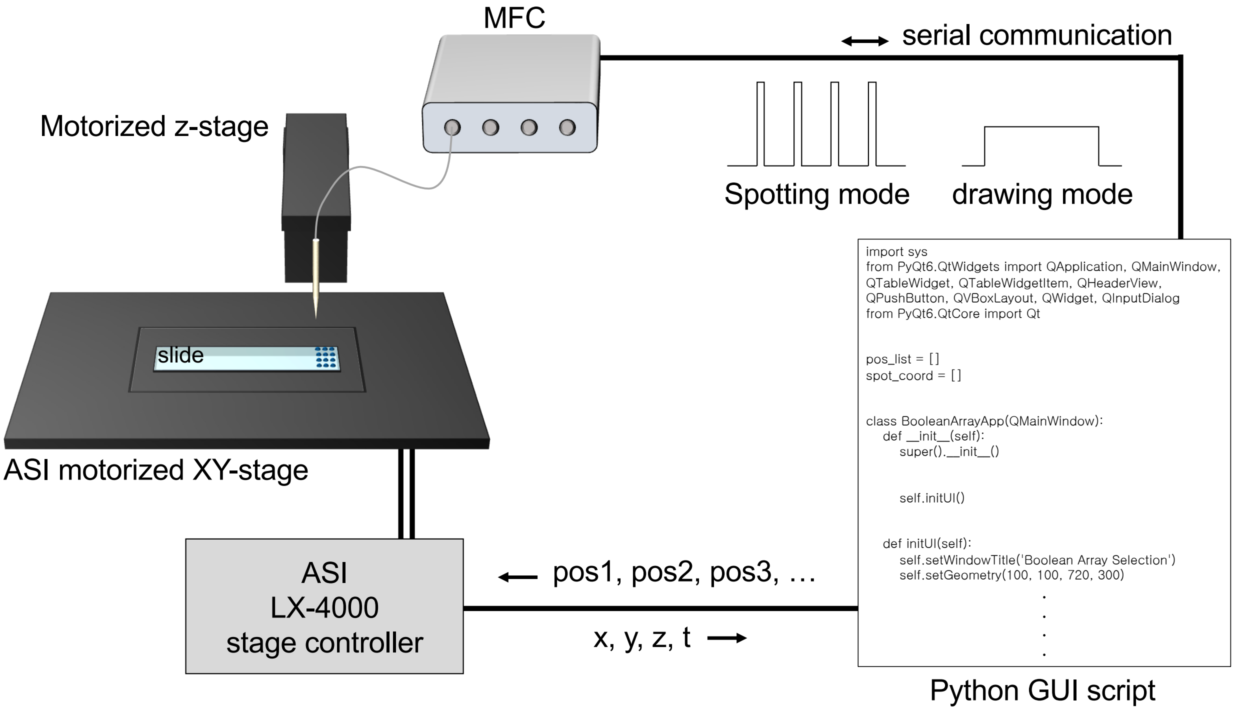


**Figure S2. Schematic diagram of automated dispenser for loading liquid reagents.** The automated dispenser is cromprised of a motorized *xy* stage, a motorized *z* stage, a stage controller, and a microfluidic flow control system (MFCS). All components are operated by a custom Python GUI script available in the OSF repository associated with the manuscript (osf.io/aqtzr).

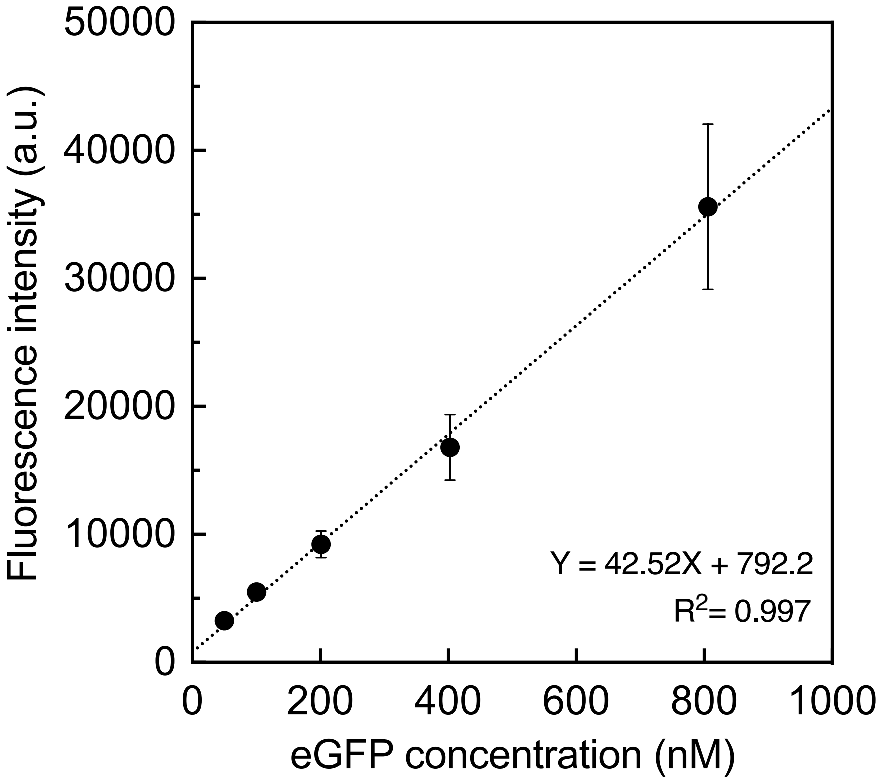


**Figure S3. Calibration curve showing measured fluorescence intensities for known eGFP concentrations.** Markers and error bars represent the median and standard deviation across 32 droplets.

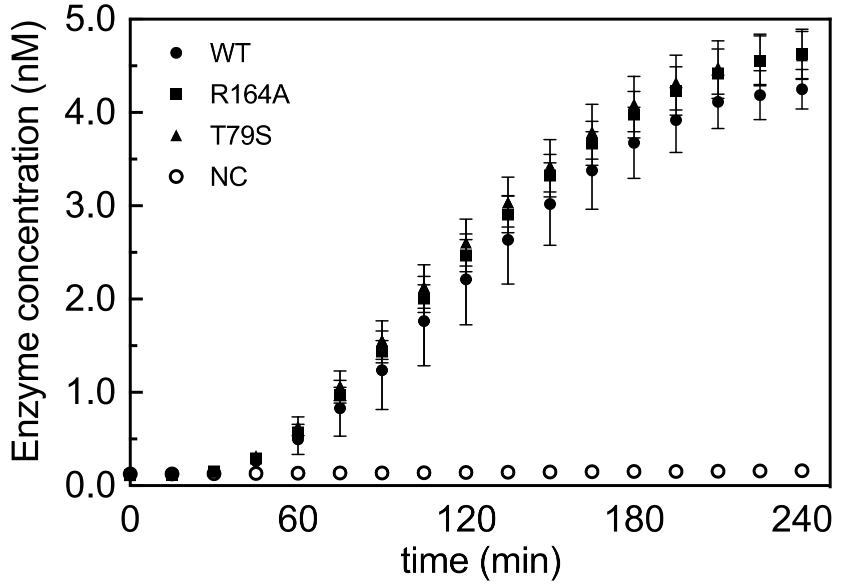


Figure S4. Expressed enzyme concentration as a function of incubation time for 3 eGFP-tagged PafA variants, WT (n = 16), R164A (n = 16), and T79S (n = 15) and a no template negative control, NC (n = 8). Markers represent the median value across all droplets and error bars represent the standard deviation of these intensities.

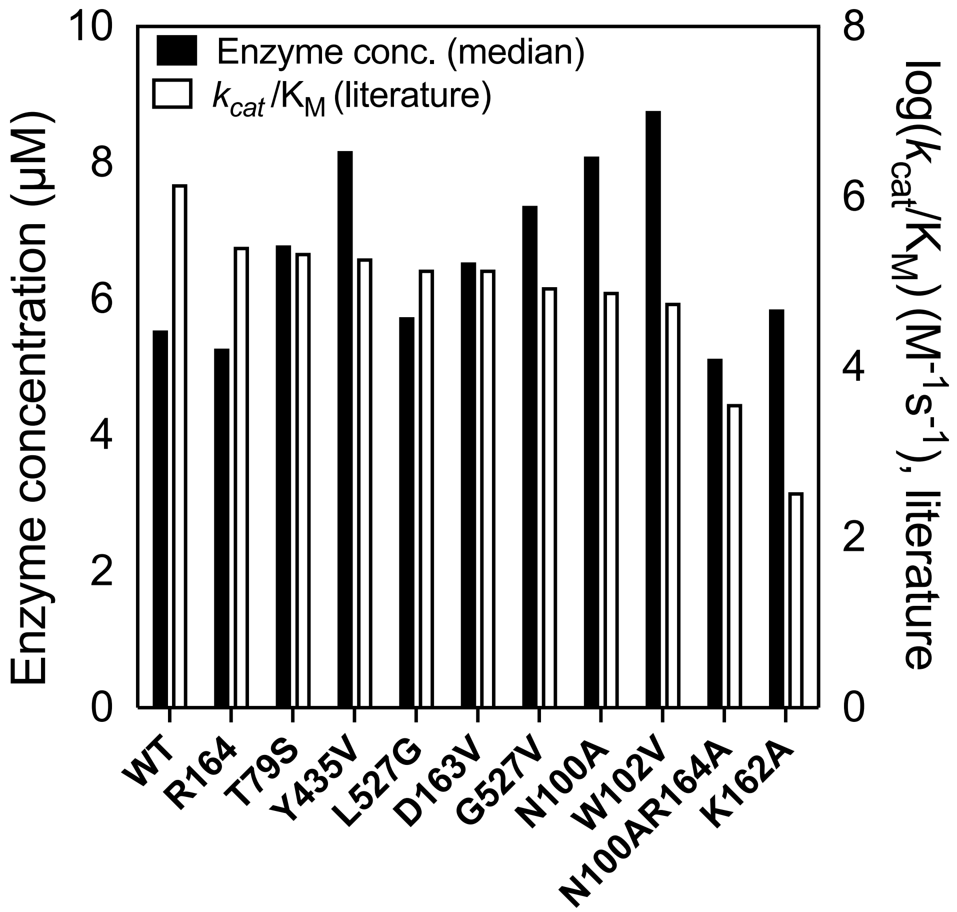


Figure S5. Comparison of median enzyme concentration (white bars, left axis; this work) and previously measured monoesterase catalytic efficiency, log($k_{cat}$/$K_M$, (black bars, right axis; literature data from Markin and Moktari *et al.*, *Science*, 2021)

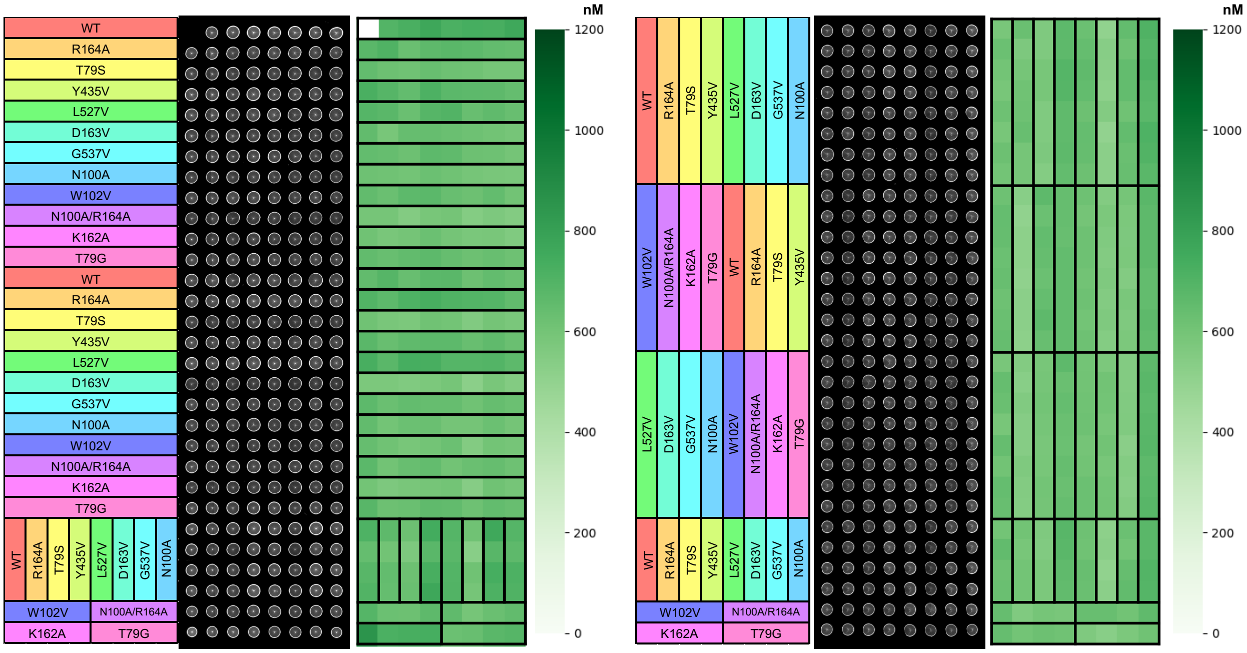


**Figure S6. Assessment of protein expression variability on the variant and positions.** Heatmaps indicating the identity of the mutant in each spot (left), observed fluorescence images (middle), and protein concentrations calculated from observed intensities (right) for two different DA-MEK slides.

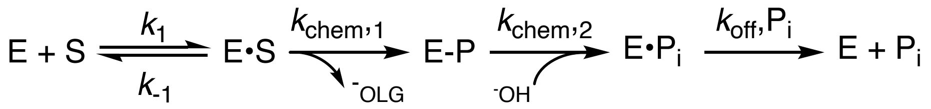


**Equation S1**. 4-MUP reaction catalyzed by PafA.

$$K_{M,obs} = K_{M,corr} \times \left(1 + \frac{[I]}{K_i}\right)$$

**Equation S2**. Correction of $K_M$ with inhibition constant ($K_i$) and inhibitor concentration ([I]).

$$V_{i,corr} = \frac{V_{max} \times [S]}{K_{M,obs}/\left(1 + \frac{[I]}{K_i}\right) + [S]}$$

**Equation S3**. Correction of initial velocities ($V_i$) at substrate concentration ([S]) with inhibition constant ($K_i$) and inhibitor concentration ([I]).

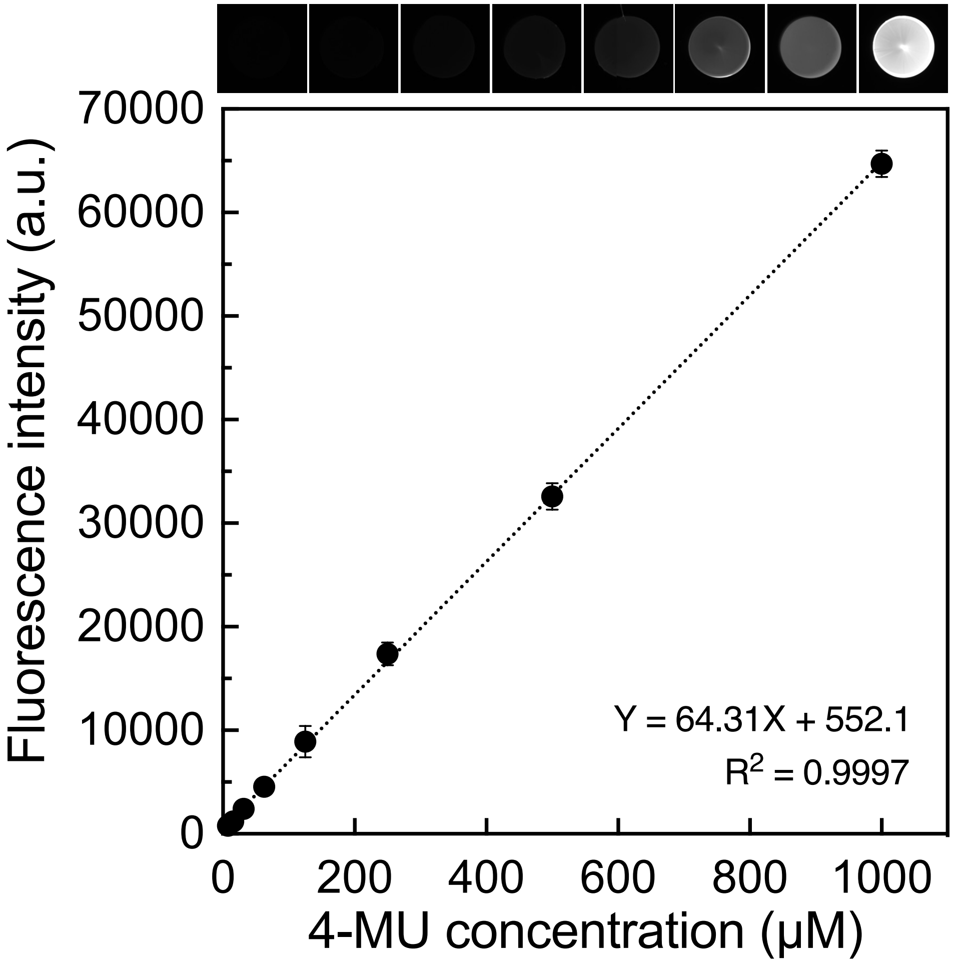


**Figure S7. Calibration curve for quantifying 4-MU fluorescence signal to concentration.** Markers indicate median intensities and error bars denote standard deviation across 16 droplets.

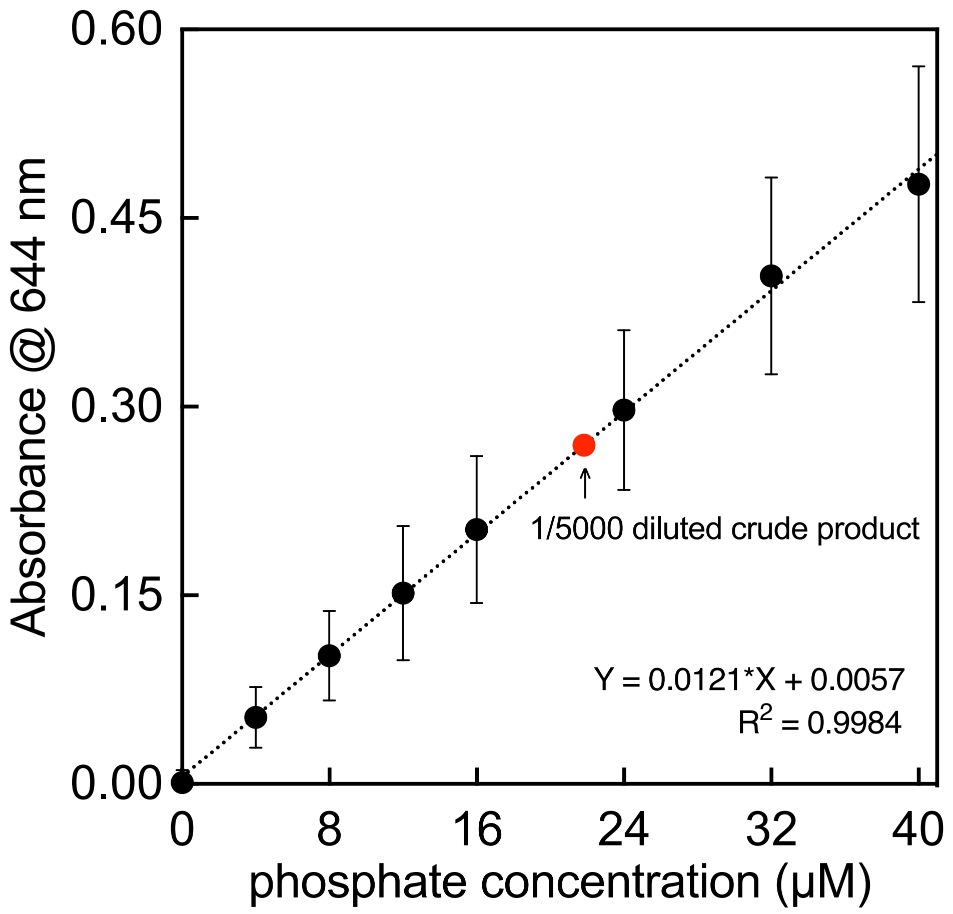


**Figure S8. Standard calibration curve relating observed absorbance to known phosphate concentration.** Black markers indicate median values and error bars indicate standard deviation across 8 wells. The red point shows an example measured value for CFPS crude product samples.

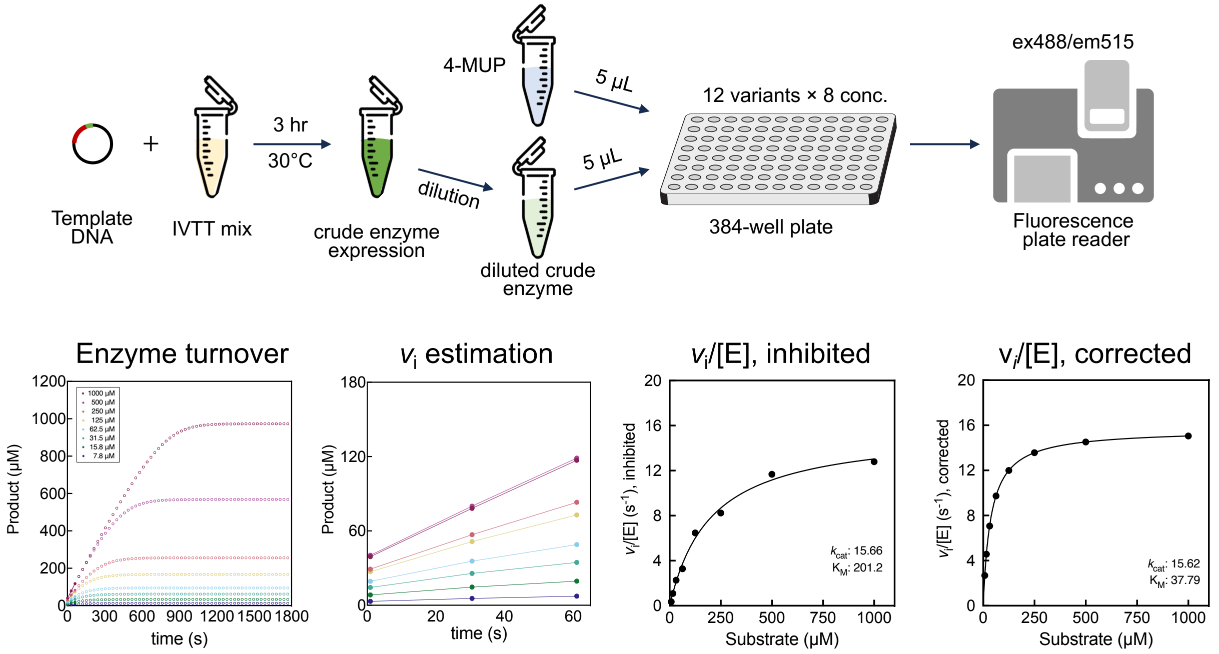


Figure S9. An experimental pipeline for well-plate-based assays measuring PafA enzyme kinetics (top row) and example measurements and fitting procedure used to calculate Michaelis-Menten constants (bottom row).

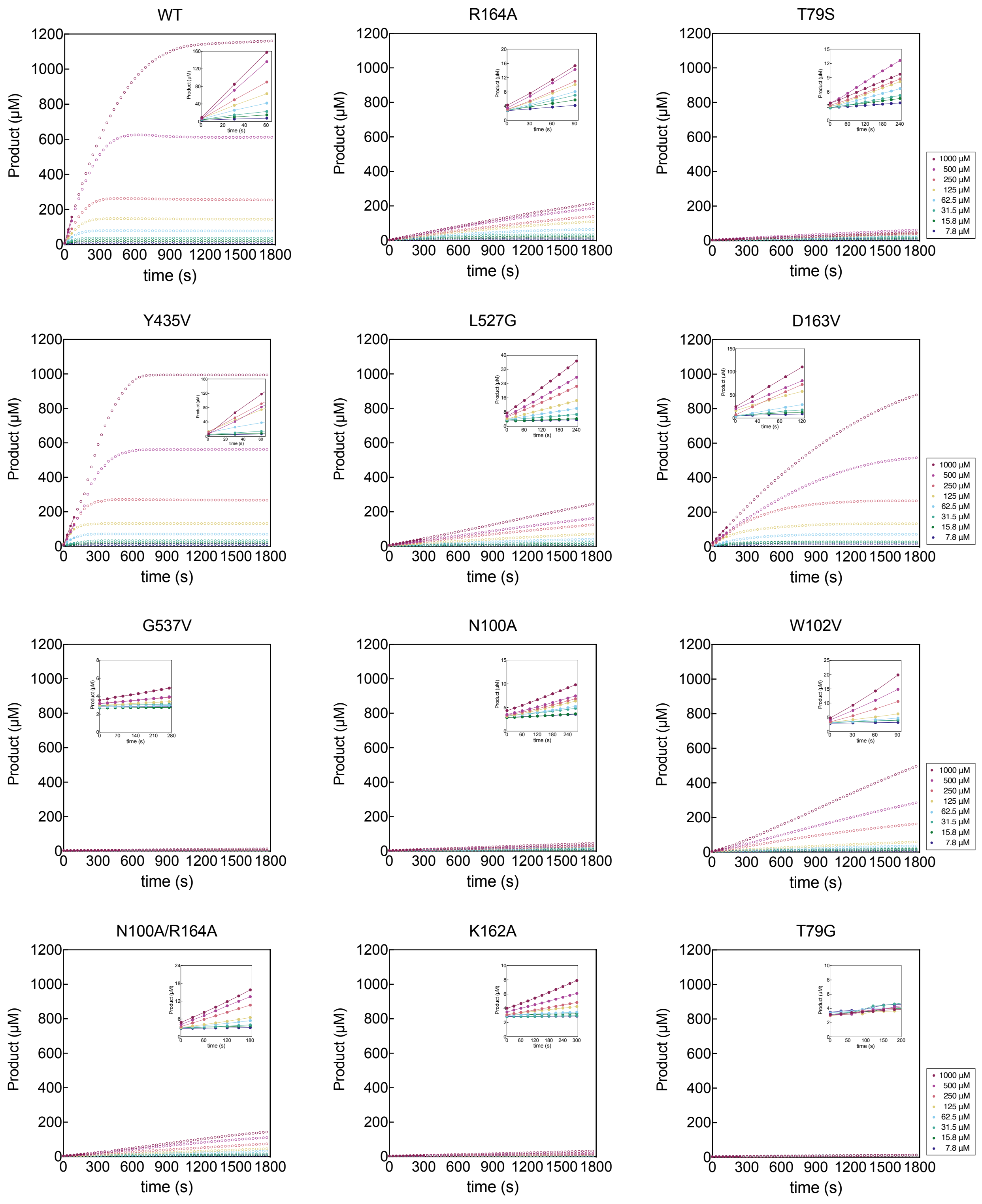


**Figure S10. Progress curves of PafA variants measured in DA-MEK.** Product (4-MU) formation was measured over time using fluorescence microscopy to determine initial velocities. The circles in each color represent the progress curve of the entire reaction, while filled circles indicate the data points included in the initial velocity calculation. These data represent single measurements corresponding to the median of all measurements for each variant, respectively.

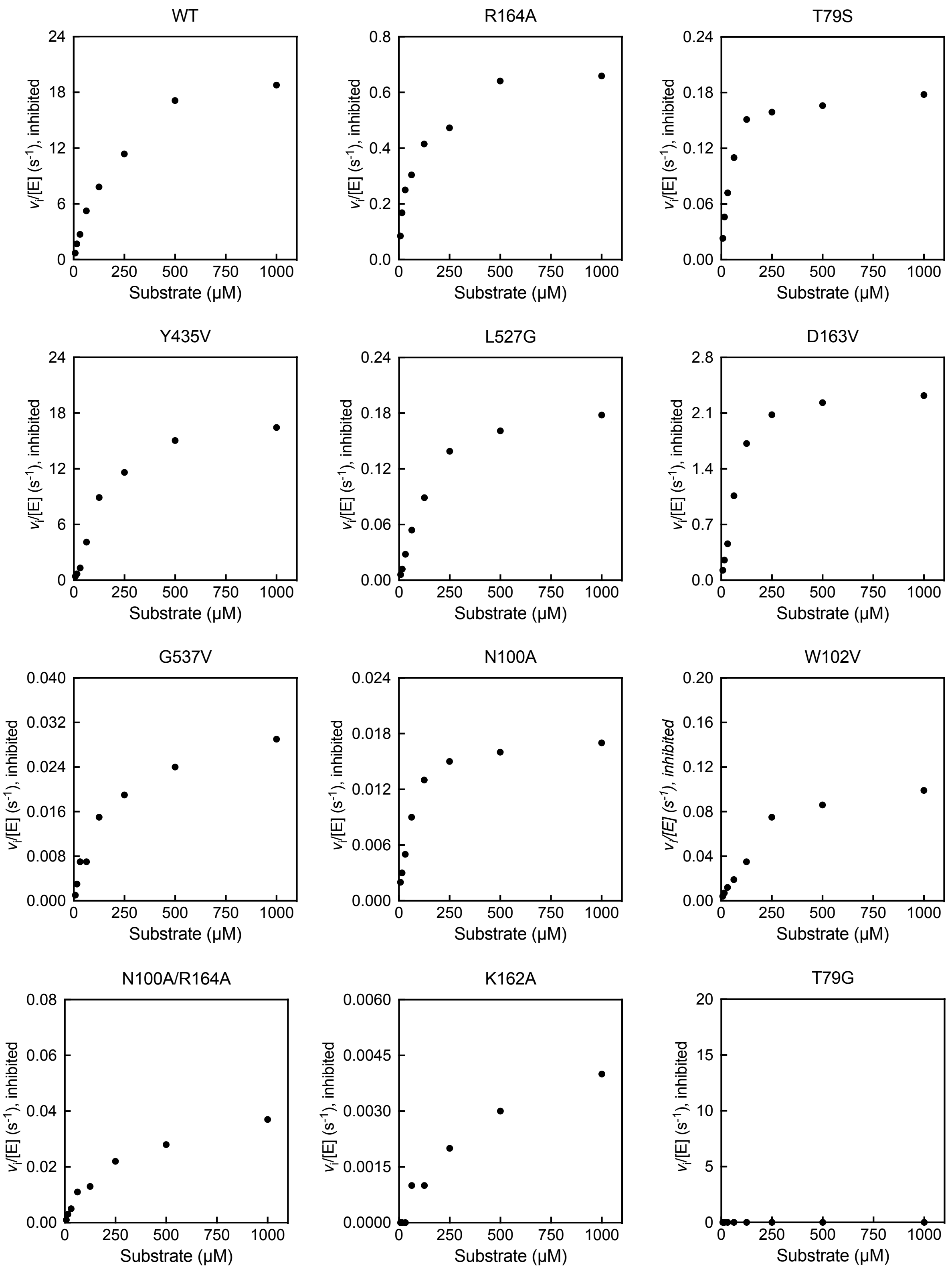


Figure S11. Uncorrected $v_i$/[E] of PafA variants observed in DA-MEK to measure $k_{cat}$ and $K_{M,\ observed}$, inhibited by phosphate content in CFPS mixtures.

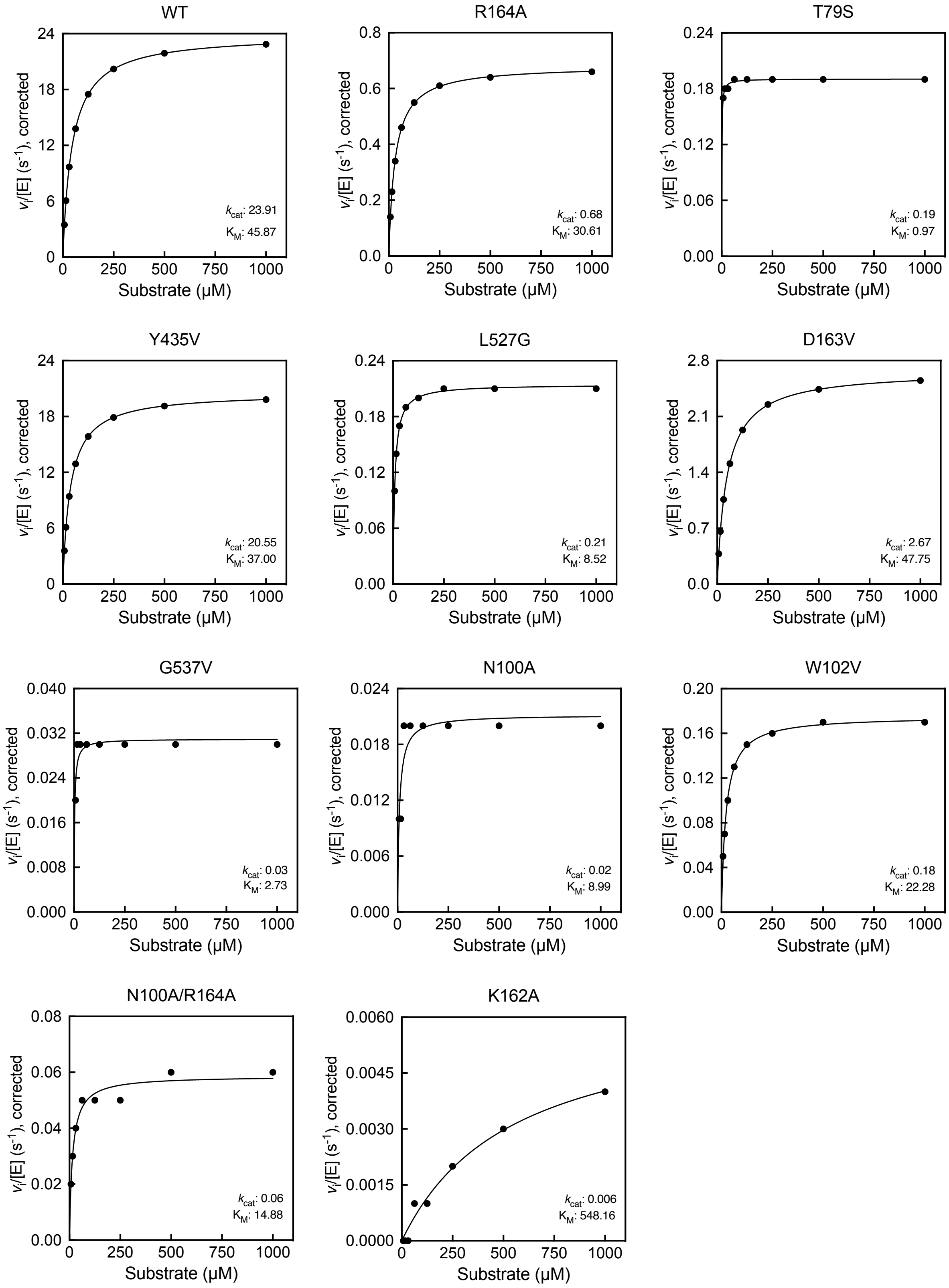


Figure S12. $v_i$/[E] of PafA variants obtained in DA-MEK, corrected for compensation of phosphate inhibition to determine $k_{cat}$ and $K_{M, corrected}$.

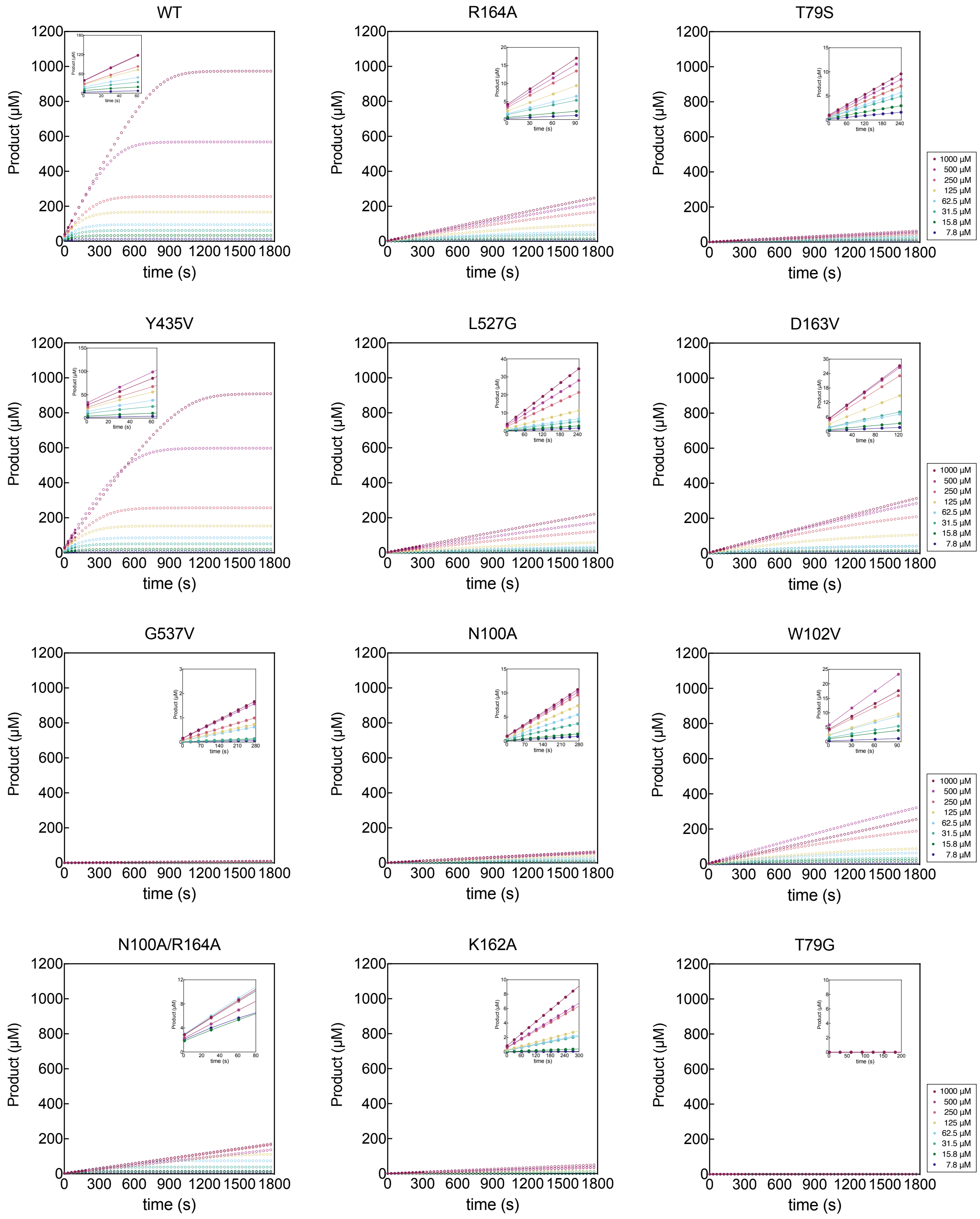


**Figure S13. Progress curves of PafA variants measured in wellplates.** Product (4-MU) formation was measured over time using fluorescence microscopy to determine initial velocities. The circles in each color represent the progress curve of the entire reaction, while filled circles indicate the data points included in the initial velocity calculation. These data represent single measurements corresponding to the median of all measurements for each variant, respectively.

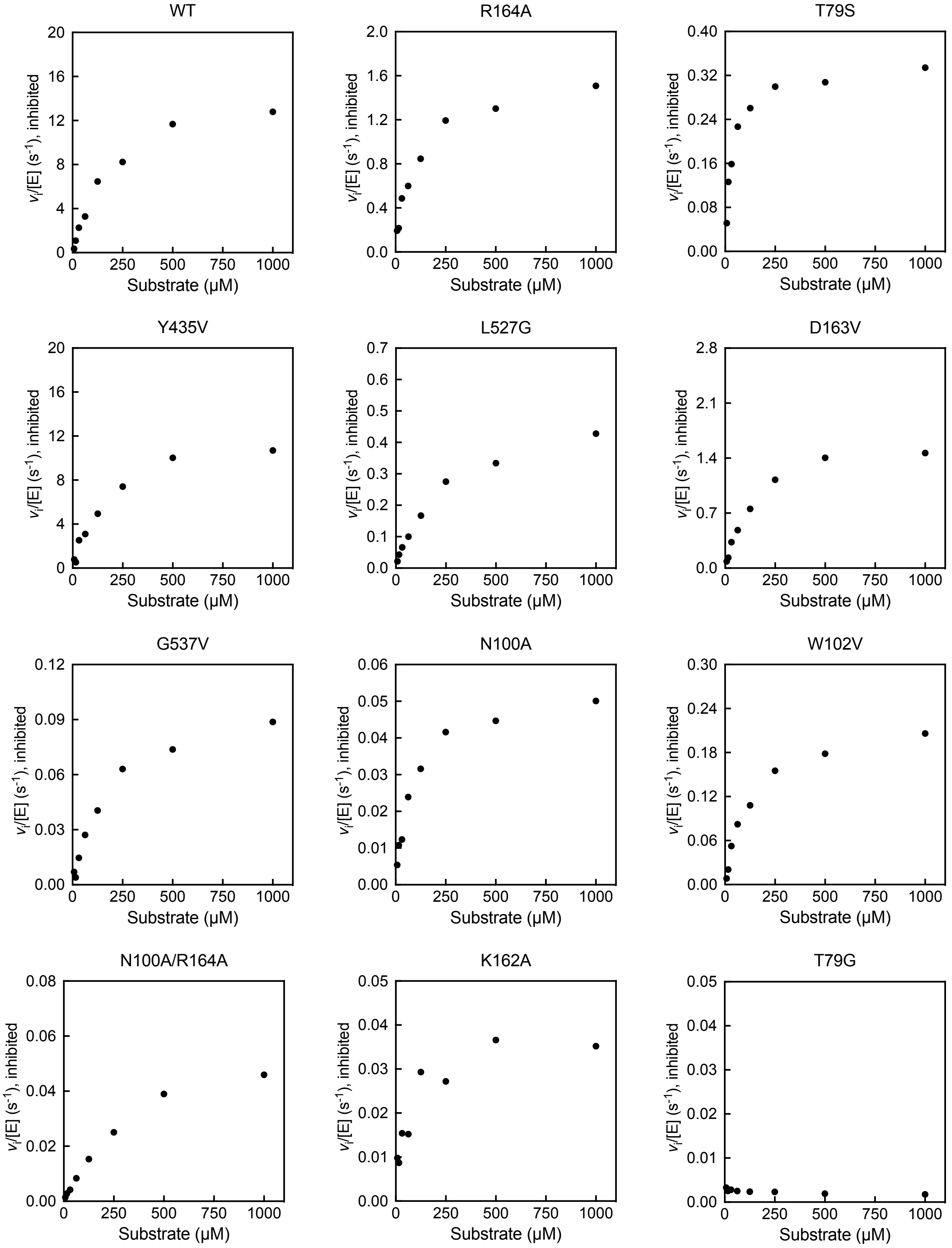


Figure S14. Uncorrected $v_i$/[E] of PafA variants observed in wellplates to measure $k_{cat}$ and $K_{M,\ observed}$, inhibited by phosphate content in CFPS mixtures

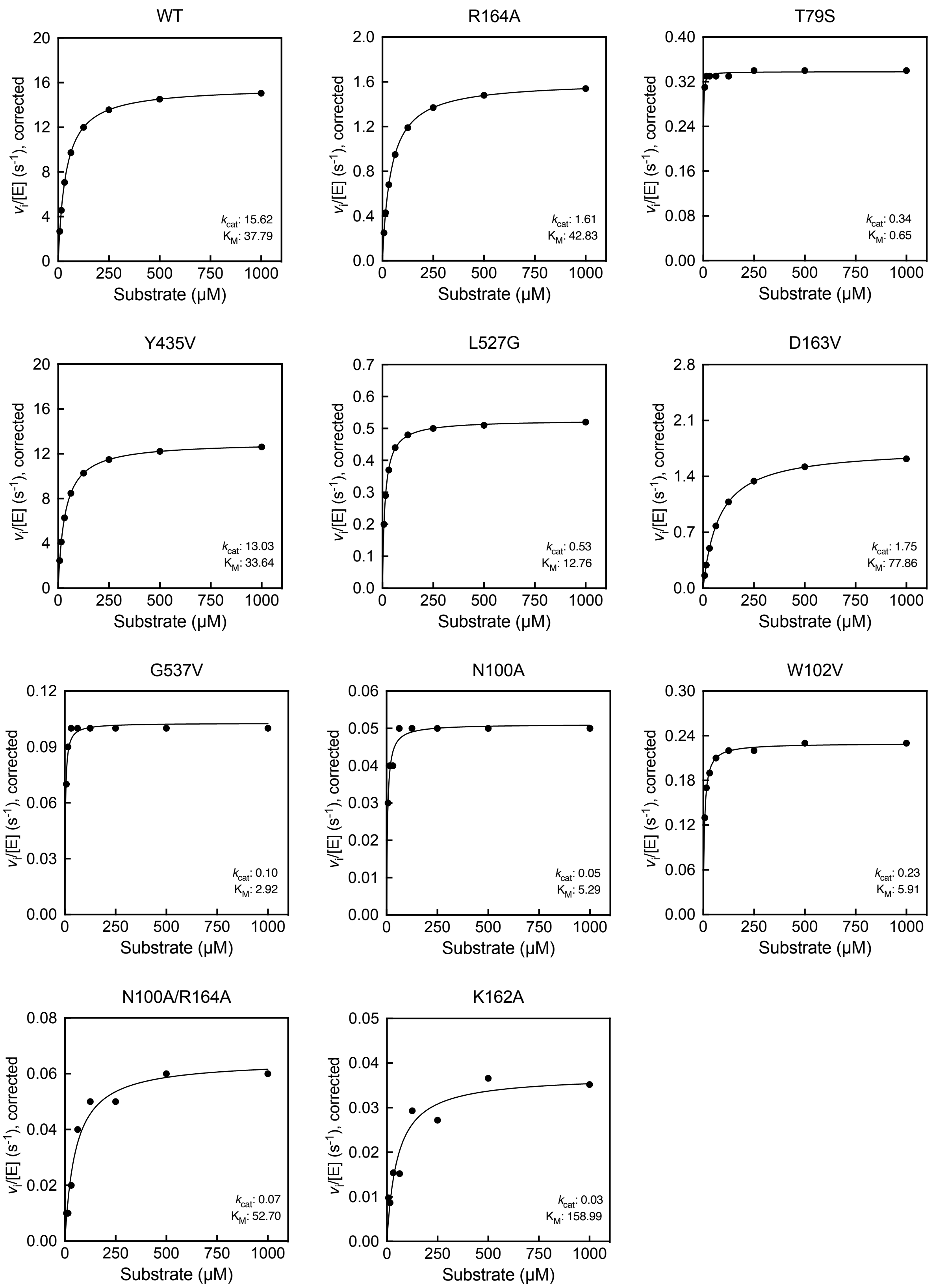


Figure S15. $v_i$/[E] of PafA variants obtained in wellplates, corrected for compensation of phosphate inhibition to determine $k_{cat}$ and $K_{M,\ corrected}$.

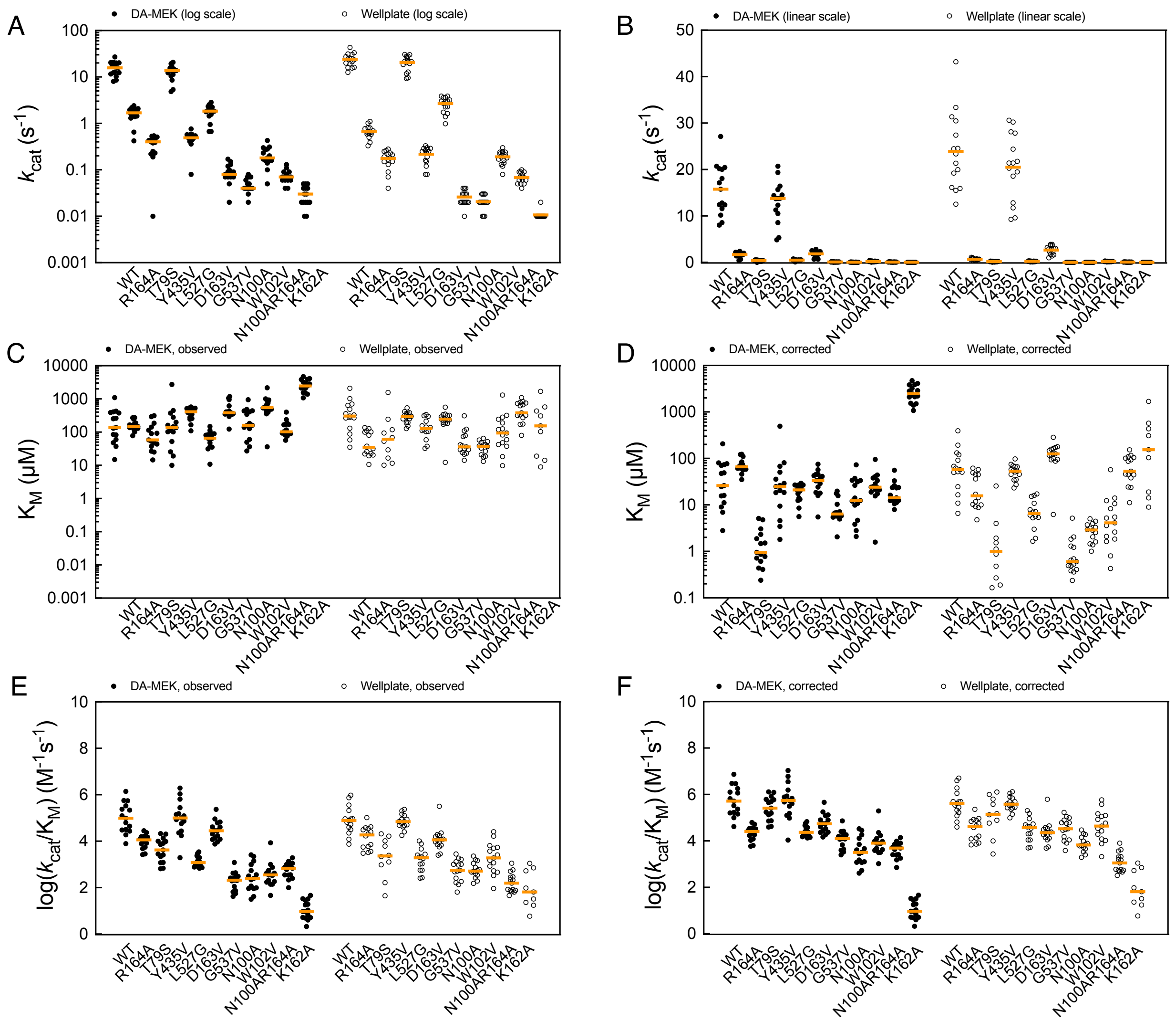


**Figure S16. Comparison of PafA variants kinetic parameters** (A) $k_{cat}$ (log scale y-axis), (B) $k_{cat}$ (linear scale y-axis), (C) observed $K_M$, (D) corrected $K_M$, (E) observed $log_{10}$-transformed $k_{cat}/K_M$, and (F) observed $log_{10}$-transformed $k_{cat}/K_M$ determined in DA-MEK (black circles) and wellplates (empty circles). Orange lines indicate the median values across the distribution. $K_M$ measurements below the lowest substrate concentration of 7.8 uM were filtered out.

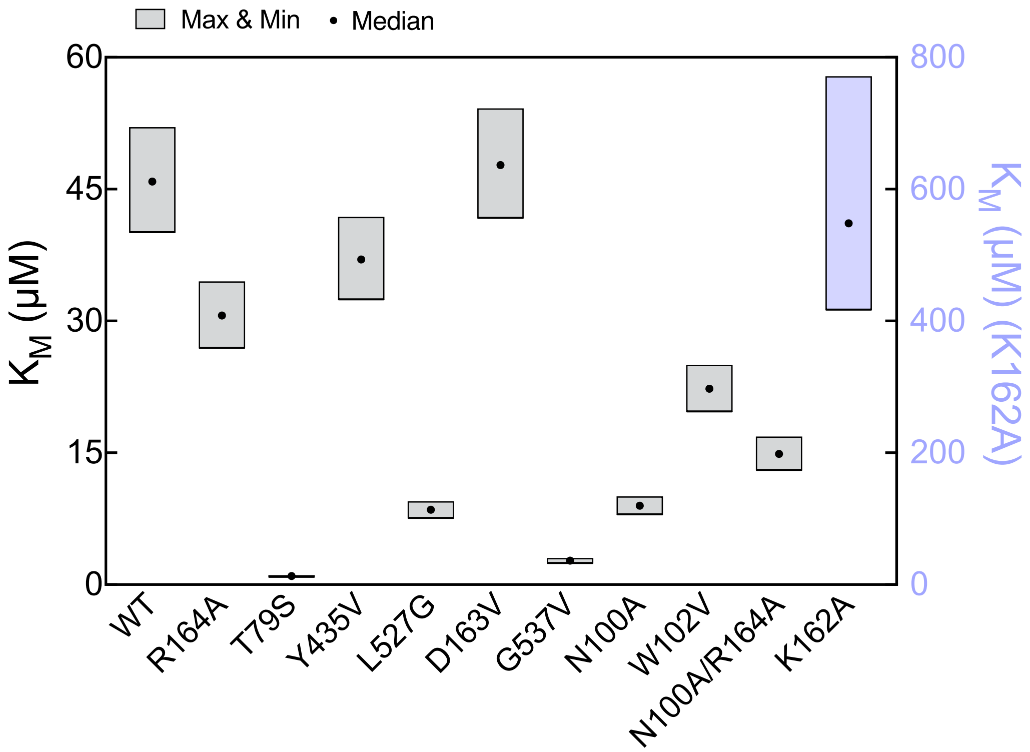


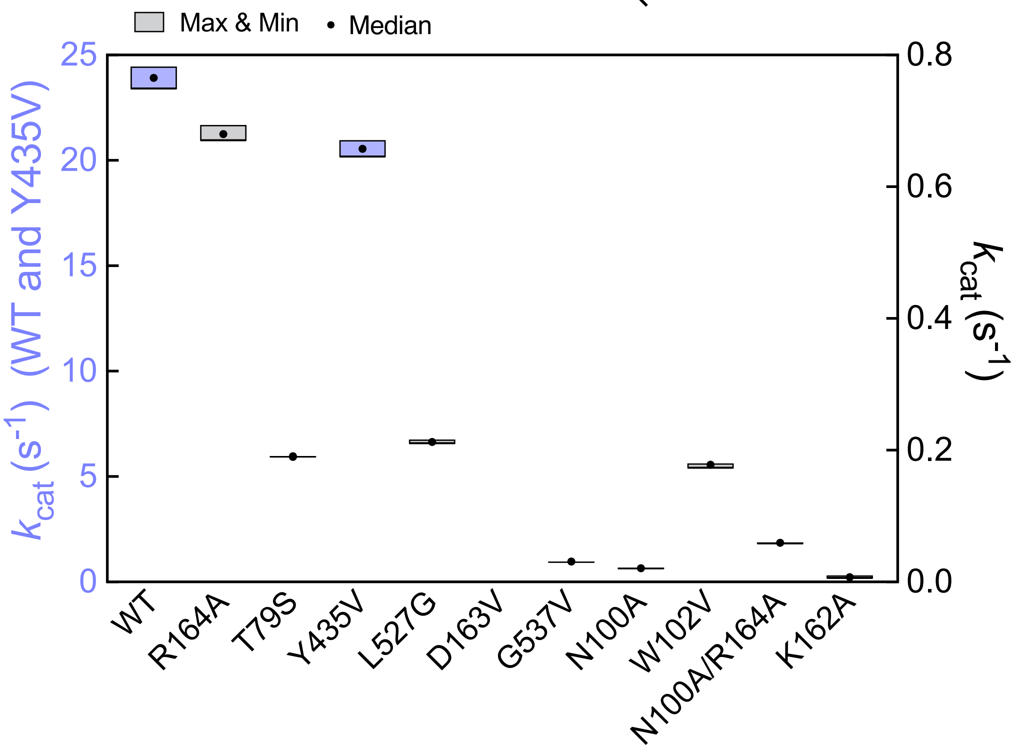


**Figure S17. Simulations to determine the range of $K_M$ and $k_{cat}$ with 10% error in substrate concentration inaccuracy.** The boxes represent the maximum and minimum values obtainable with a 10% error, while the black dots represent the median of actual measurements.

**Table S1. Comparison of costs for screening a variant library using DA-MEK or 384-well plates.** The required volumes for each reaction are half of the total reaction volumes.

| | DA-MEK | | | Well plate |
|---|---|---|---|---|
| Type | 500 μm (dia.) | 1000 μm (dia.) | 1500 μm (dia.) | 384-well |
| Volume/rxn (μL) | 0.02 | 0.13 | 0.45 | 5.00 |
| Cost/volume ($/μL) | 1.18 | 1.18 | 1.18 | 1.18 |
| Cost/rxn ($) | 0.02 | 0.15 | 0.53 | 5.90 |
| Number of conc. | 8 | 8 | 8 | 8 |
| Replicates/conc. | 3 | 3 | 3 | 3 |
| Cost/variant ($) | 0.57 | 3.68 | 12.74 | 141.60 |
| Library size | 1,000 | 1,000 | 1,000 | 1,000 |
| Total cost ($) | $566 | $3,682 | $12,744 | $141,600 |
| Ratio | 1 | 6.5 | 22.5 | 250.0 |
